# Individualized prediction of future cognition using baseline developmental changes in cortical anatomy

**DOI:** 10.1101/2021.07.05.451172

**Authors:** Budhachandra Khundrakpam, Linda Booij, Seun Jeon, Jussi Tohka, Alan C. Evans

## Abstract

Predictive modeling studies have started to reveal brain measures underlying cognition; however, most studies are based on cross-sectional data (static ‘final’ brain measures acquired at one time point). Since brain development comprises of continuously ongoing events leading to cognitive development, predictive modeling studies need to consider *‘dynamic’* as opposed to *static ‘final’* brain measures. Using longitudinal neuroimaging and cognitive data (global executive composite score, an index of executive function) from 82 individuals (aged 5-14 years, scanned 3 times), we built highly accurate prediction models (*r*=0.61, *p*=1.6e-09) of future cognition (assessed at visit 3) based on baseline developmental changes in cortical anatomy (from visit 1 to 2). More importantly, dynamic brain measures (change in cortical anatomy from visit 1 to 2) and not static brain measures (cortical anatomy at visit 1 and 2) were critical for predicting future cognition, suggesting the need for considering dynamic brain measures in predicting cognitive outcomes.

## Introduction

Understanding the neurobiology of cognition is the central tenet of cognitive neuroscience. Much of the previous efforts, however, have focused primarily on group-level studies that utilized univariate methods with structural and functional brain measures. Broadly, these studies have either focused on *i*) finding brain measures with significant group differences between healthy individuals and individuals with brain disorders using statistical inferences, and *ii*) determining relation between brain measures and behavior using correlational analysis. Although group-level studies provided important insights into the human brain, there may be considerable heterogeneity in brain measures and behavior within the group (1–4), information critical for understanding the neurobiology of cognition (5, 6). Further, univariate methods usually focus on finding cross-sectional associations between observed behavior and brain measures, instead of predicting behavior using brain measures (7).

Predictive modeling, on the other hand, utilizes machine-learning methods to assess individual variability in cognition using brain measures (5, 8–12). More specifically, predictive modeling has been used to achieve predictions of various cognitive measures, such as intelligence (13–16), attention (6), working memory (17–19), reading comprehension (20), inhibition control (12, 21), cognitive flexibility (12, 22, 23) and creativity (24, 25) using neuroimaging features of functional or structural connectivity, gray matter volume, cortical thickness, and fractional anisotropy.

Much of the predictive modeling studies so far are based on data that acquire brain measures and cognitive measures at the same time. Hence, the developmental time course of the associations remains unknown. However, the real scope of predictive modeling lies in predicting long-term cognitive outcomes based on baseline neuroimaging data (26, 27). A shortcoming in these studies is that they used *static ‘final’* brain measures (read as brain measures acquired at one time point) to predict cognitive outcomes. Brain development is a continuously ongoing process leading to cognitive development. In fact, studies have shown the role of cortical trajectories in cognitive development (28). There is thus a need for considering *trajectories* as opposed to *static ‘final’* brain measures. We therefore need to build prediction models which incorporate developmental changes in brain measures.

Thus, the aim of our study was to systematically compare the utility of static and dynamic brain measures in the prediction of cognitive outcome. To this end, we utilized longitudinal neuroimaging data, collected at 3 time points from individuals who also had concurrent measures of cognitive function. Specifically, in the present analyses, within-subject developmental changes in brain morphometry (computed as the change in cortical thickness from time point 1 to 2) was used to build prediction models of executive function at time point 3. We chose executive function (EF) as our measure of cognition because EF is a vital cognitive domain which is adversely affected in several neurodevelopmental and psychiatric conditions (29). We hypothesized that incorporating within-subject developmental changes in brain measures will yield better predictive models of cognition compared to those with ‘static’ (read as, crosssectional) brain measures.

## Methods

### Subjects

Longitudinal MRI scans were taken from the NIH MRI Study of Normal Brain Development (NIHPD) repository (30). The study sample was similar to our previous publication (31) and consisted of participants that were scanned at three separate times approximately 2 years apart (*n* = 82, age (mean ± standard deviation) = 11.9 ± 2.9 years, males/females = 31/51, **Figure 1**). In terms of separate visits, the age of participants was 9.9 ± 2.4 years for visit 1, 11.9 ± 2.5 years for visit 2 and 13.8 ± 2.4 years for visit 3.

**Figure 1.**
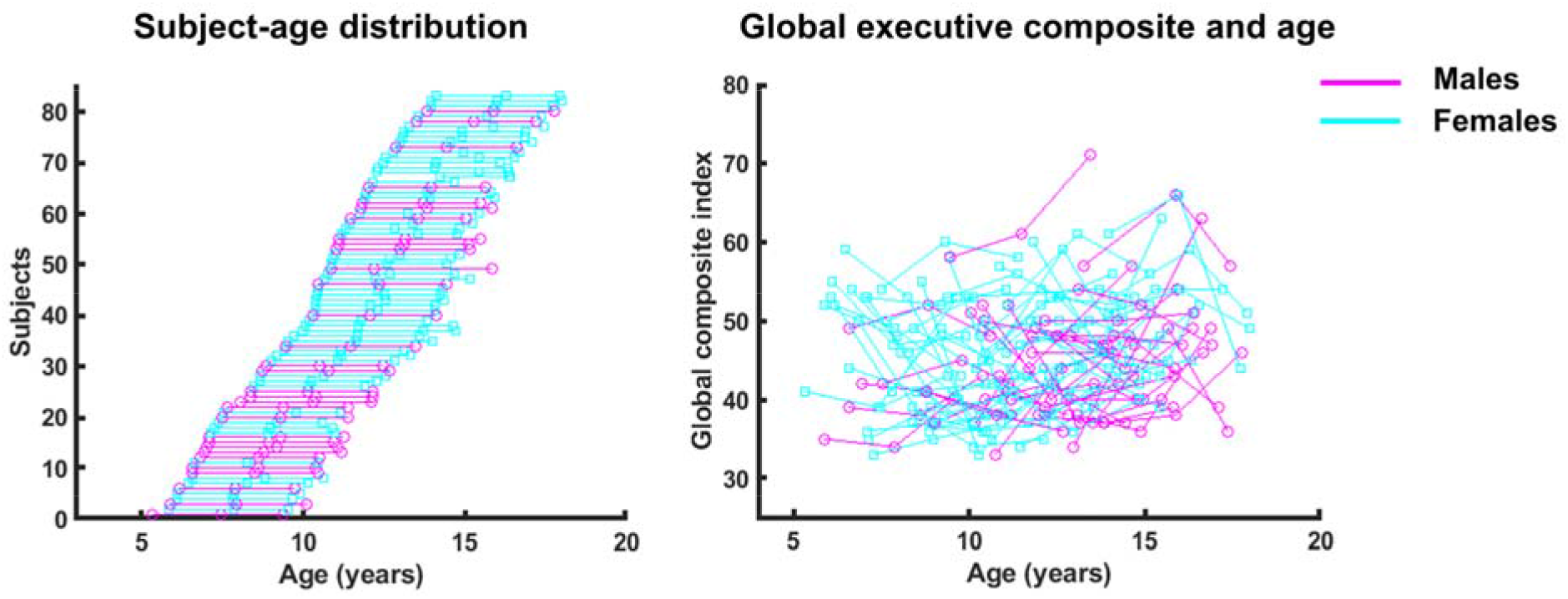
Subject-age distribution and distribution of Global executive composite with age. Participants were scanned at ~ 2 years apart. Global executive composite (GEC) was age standardized.

### Cognitive measure

For cognitive measure, we used the Global executive composite (GEC) available with the Behavior Rating Inventory of Executive Functions (BRIEF) (32). The BRIEF is a well-known and well-validated measure assessing dimensions of executive function in children and adolescents as manifested in everyday life. The parent version of the questionnaire was administered. The BRIEF composes of three summary indices: Behavioral Regulation, Meta-cognition, and an overall score, the Global Executive Composite. *T*-scores are generated for each index. For the current analyses, we focused only on the overall index – the Global executive composite, henceforth referred as GEC. In terms of visits, the GEC scores ranged from 33 to 61 (mean ± standard deviation = 45.9 ± 7.2) for visit 1, 33 to 66 (45.2 ± 7.7) for visit 2 and 35 to 71 (46.2 ± 7.1). The scores were age-standardized, and hence did not change significantly between the three visits.

### Image pre-processing

The CIVET processing pipeline, (http://www.bic.mni.mcgill.ca/ServicesSoftware/CIVET) developed at the Montreal Neurological Institute, was used to compute cortical thickness measurements at 81,924 regions covering the entire cortex. A summary of the steps involved follows; the *T_1_*-weighted image is first non-uniformity corrected, and then linearly registered to the Talairach-like MNI152 template (established with the ICBM152 dataset). The non-uniformity correction is then repeated using the template mask. The non-linear registration from the resultant volume to the MNI152 template is then computed, and the transform used to provide priors to segment the image into GM, WM, and cerebrospinal fluid. Inner and outer GM surfaces are then extracted using the Constrained Laplacian-based Automated Segmentation with Proximities (CLASP) algorithm, and cortical thickness is measured in native space using the linked distance between the two surfaces at 81,924 vertices. Each subject’s cortical thickness map was blurred using a 30-millimeter full width at half maximum surface-based diffusion smoothing kernel to impose a normal distribution on the corticometric data, and to increase the signal to noise ratio. Quality control (QC) of the data was performed by two independent reviewers, and only scans with consensus of the two reviewers were used. As a result of this process, data with motion artifacts, a low signal to noise ratio, artifacts due to hyperintensities from blood vessels, surface-surface intersections, or poor placement of the grey or white matter (GM and WM) surface for any reason were excluded.

### Univariate analysis

The association of static brain measures (cortical thickness at visit 1 and 2) and BRIEF scores at visit 3 was analyzed using a vertex-wise general linear model (GLM) and was quantified using *t*-statistics. Significant association was assessed with multiple comparison corrected *p*-statistics using random field theory (RFT) (33). Similar analysis was performed for the association of dynamic brain measures (slope of the change in cortical thickness from visit 1 to 2) and BRIEF scores at visit 3. Age and sex were included as covariates. GLM analysis was performed using SurfStat (http://www.math.mcgill.ca/keith/surfstat/).

### Predictive model

BRIEF scores were predicted based on Random Forest regression analysis (34). Random (regression) forests are an ensemble supervised learning method for regression that operate by constructing a multitude of regression trees at training time and outputting the mean prediction of the individual trees for given input. Regression trees are constructed utilizing bagging, random sampling (with replacement) of the training set to avoid overfitting to the training set. We utilized the *TreeBagger* implementation of Matlab (version 2019b) and set the number of trees in the ensemble to 1000. This number of trees provided clear convergence of the out-of-bag error (34, 35). Other parameters of the random forests were set to their default values in the *TreeBagger* implementation; in particular, the *mtry*-parameter was set to *d*/3, where *d* is the number of predictor variables.

Random forests were trained with different sets of predictor variables based on cortical thickness and behavioral data. However, all the predictor variables were from the time points *t_1_* and *t_2_* to predict the behavior at *t_3_*. Cortical thickness was processed into slopes of cortical thickness as presented in **Figure 2**. Thereafter, we regressed out the age at *t_1_*, square of age at *t_1_*, gender as well as (gender × age) interaction. The so-formed residuals were *z*-transformed, and then entered into a principal component analysis. The scores of the principal components that explained more than 1.5% of variance were then used as cortical thickness predictors. The cortical thickness values at *t_1_* and *t_2_* went through the same processing, only the number of principal components was constrained to be the same (number of principal components = 14) as with the CT slopes.

**Figure 2.**
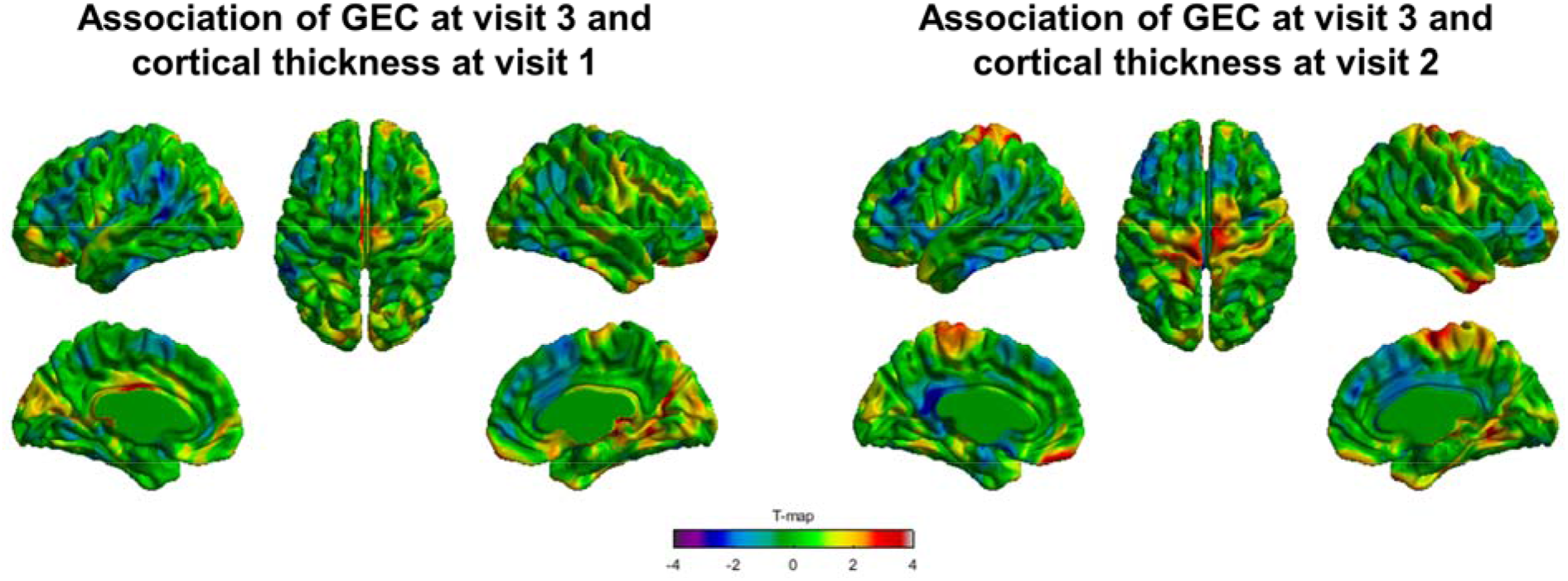
Correlation analysis of cognitive measure at visit 3 and static brain measures. *T*-maps of association between GEC at visit 3 and cortical thickness at visit 1 (image on the left) and GEC at visit 3 and cortical thickness at visit 2 (image on the right). None of the brain regions showed significant association after correction for multiple comparisons using random field theory, RFT. Note, GEC = Global executive composite.

We estimated the generalization error for the predictions via out-of-bag error. As explained more in detail in (34), for a sample (*x,y*), out-of-bag predictor aggregated predictions of those regression trees which were not trained using (*x,y*) due to bagging. Then, the prediction error was computed based on the out-of-bag predictions. These out-of-bag errors was computed at the training time, without the need for re-training as in cross-validation. It is to be noted that, out-of-bag error estimates are equally or more accurate than the cross-validation or holdout error estimates (34–36). Moreover, as the out-of-bag prediction was done for each individual sample, this enabled us to use the permutation test developed in (37) to compare the regression models as well as to compute the mean absolute error and mean square error.

## Results

For the study, we used longitudinal MRI of 82 individuals with age (mean ± standard deviation) = 11.9 ± 2.9 years, males/females = 31/51, **Figure 1**) who were scanned at three separate times approximately 2 years apart, taken from the NIH MRI Study of Normal Brain Development (NIHPD) repository (30).

1. ***Correlation analysis of cognition at visit 3 using static brain measures (cortical thickness at visit 1 and 2):*** We first investigated correlation analyses using GLM to study the association between static brain measures (cortical thickness at visit 1 and 2) and GEC BRIEF scores at visit 3. We did not observe any significant association between GEC at visit 3 and cortical thickness at visit 1 (**Figure 2A**) and visit 2 (**Figure 2B**). However, trend-level association was observed between GEC at visit 3 and cortical thickness at visit 2 in brain regions localized in sensorymotor cortices (**Figure 2B**).
2. ***Correlation analysis of cognition at visit 3 using dynamic brain measures (slope of cortical thickness from visit 1 to 2):*** We observed significant association between GEC at visit 3 and dynamic brain measures (slope of the change in cortical thickness from visit 1 to 2) (**Figure 3A**). A significant association (random field theory, RFT-corrected for multiple comparisons) was observed at one peak localized in the left-hemispheric postcentral gyrus. At this peak, the slope of the change in cortical thickness was significantly associated (*T* = 4.57, *p* < 0.01) with GEC at visit 3 (**Figure 3B**).
3. ***Prediction of cognition at visit 3 using dynamic brain measures:*** We next performed random forest regression analysis for prediction of future cognition (**Figure 5**). We observed a significant correlation of *r* = 0.61, *p* = 1.6e-09 between the true and predicted GEC (at visit 3) scores (**Figure 6A**). Additionally, our analysis using principal components of the data input based on out-of-bag (OOB) permutation variable importance (34) revealed clusters localized in the prefrontal and parietal cortices as the top model predictors (**Figure 6B**).
4. ***Comparative analysis of predictive models of cognition at visit 3 using static and dynamic brain measures:*** Hypothesis testing was performed with respect to the predictions of the random forest (RF) model using GEC_t1_ and GEC_t2_. These tests were one-sided and were used to confirm whether a prediction model performed better than our baseline model RF model (GEC_t1_ + GEC_t2_). Using demographic data (age and sex) as input, the prediction model performed badly (**Table 1**), confirming that GEC_t3_ contained no age-nor sex-related bias. Interestingly, we observed that the prediction models also performed badly if only brain measures (and no cognitive measures from visit 1 and 2) were used as input (**Figure 7, Table 1**). Using static brain measures (Thick_t1_, Thick_t2_, Thick_t1_ + Thick_t2_) as input in addition to GEC_t1_ and GEC_t2_ improved the prediction over the baseline, e.g., MSE dropped from 39.07 to 36.69, but the improvement did not reach significance (*p* = 0.193). When we used dynamic brain measures (Slope), we observed that MSE dropped from 39.07 to 32.97. This improvement in MSE was significant (*p* = 0.028, one-sided) for RF model (GEC_t1_ + GEC_t2_ + Slope) compared to RF model (GEC_t1_ + GEC_t2_).

**Figure 3.**
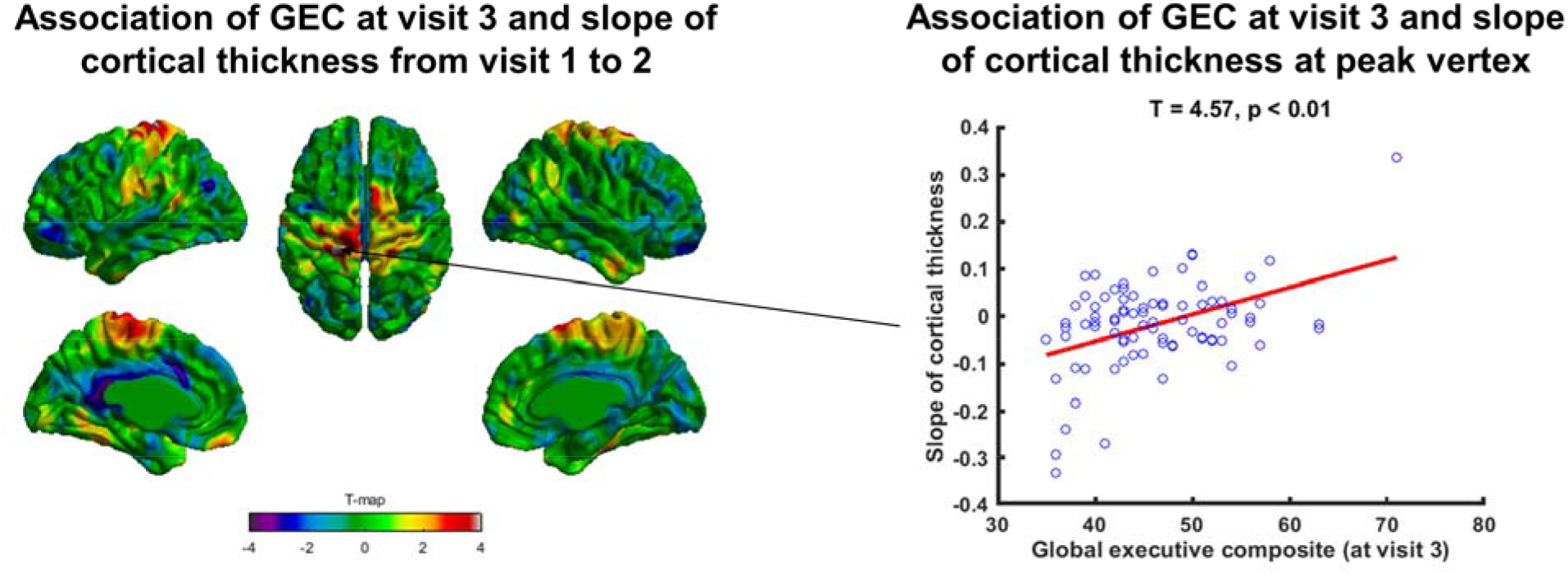
Correlation analysis of cognitive measure at visit 3 and dynamic brain measures. *T*-map of association between GEC at visit 3 and slope of change in cortical thickness from visit 1 to 2 is shown on the left side. Significant association (RFT-corrected for multiple comparisons) was observed at one peak localized in the left-hemispheric postcentral gyrus. At this peak, the scatter plot for slope of change in cortical thickness vs GEC at visit 3 was plotted showing significant association (*T* = 4.57, *p* < 0.01). Note, GEC = Global executive composite, RFT = random field theory.

**Figure 5.**
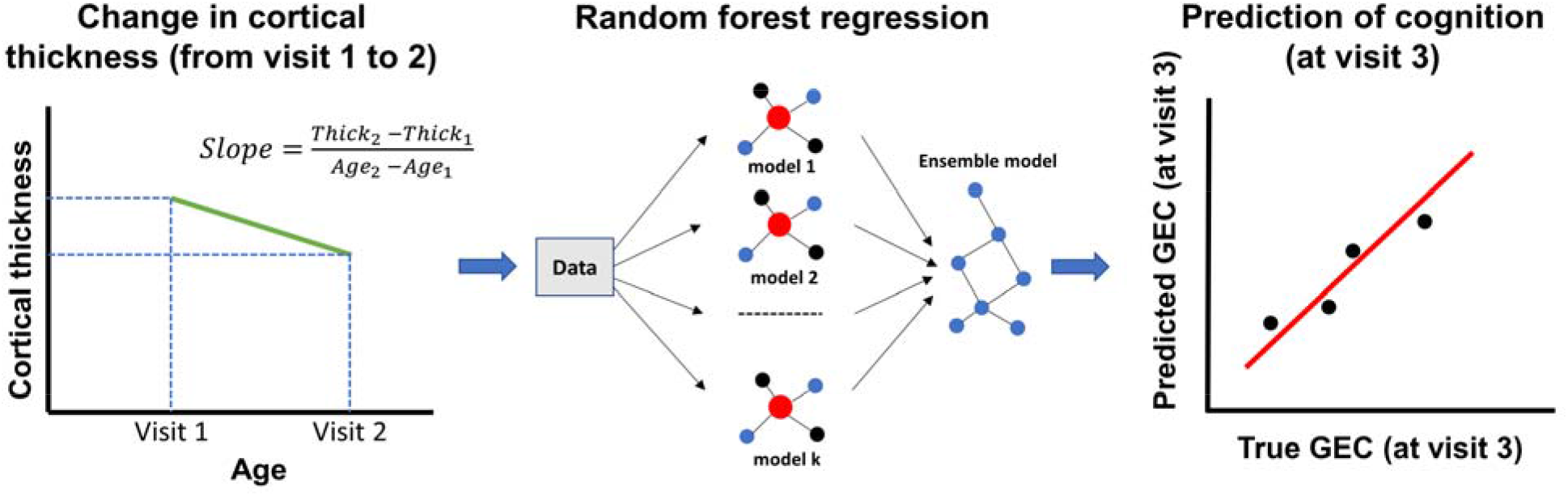
Schematic of the model for predicting cognition based on dynamic brain measures. Input data for the prediction model is dynamic brain measure, computed as the slope of the change in cortical thickness from visit 1 to 2. The prediction model is random forest regression model. The cognitive measure for prediction is the Global executive composite at visit 3. GEC = Global executive composite.

**Figure 6.**
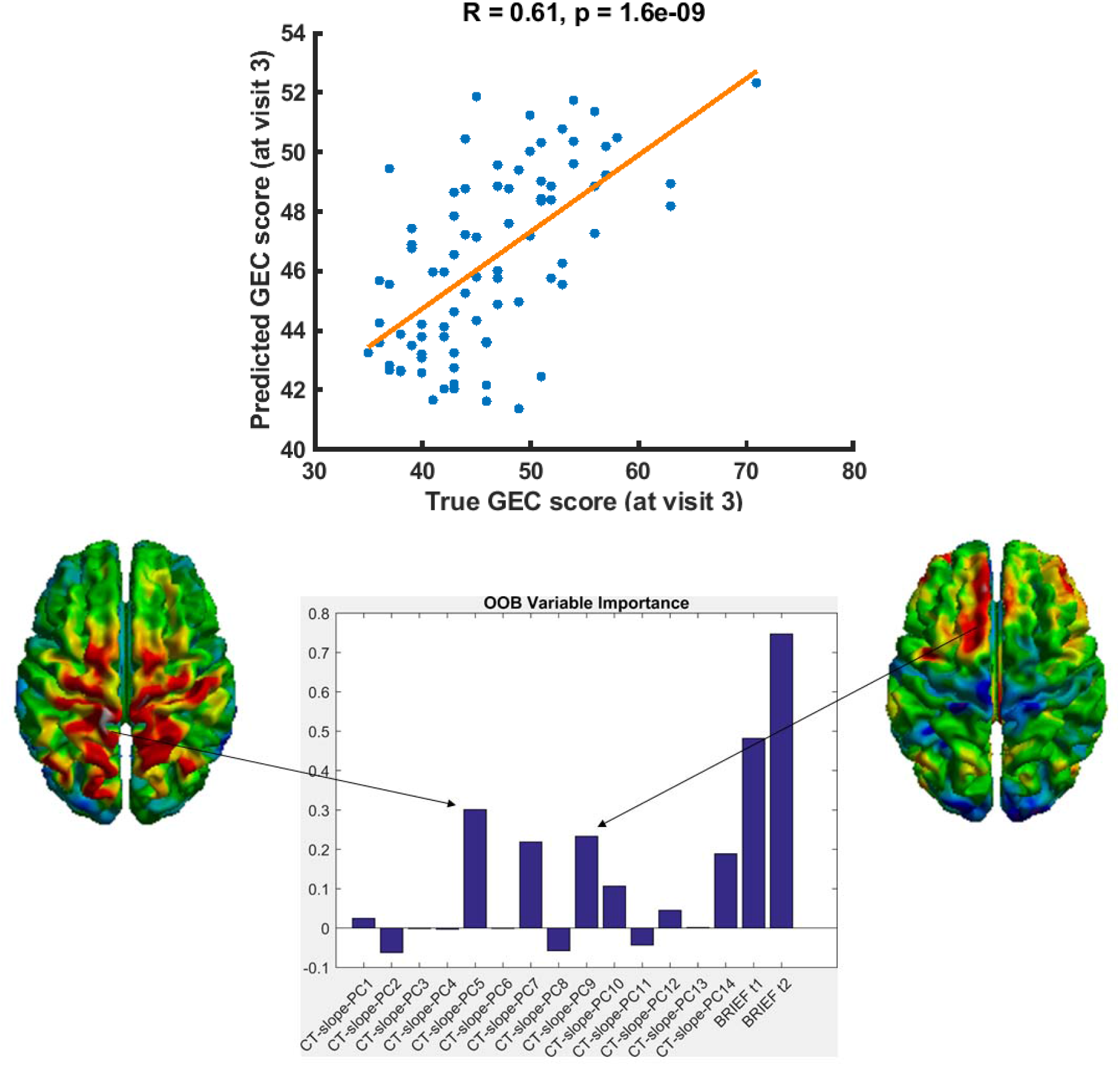
Prediction of cognitive measure at visit 3 using dynamic brain measures. Using dynamic brain measures (slope of the change in cortical thickness from visit 1 to 2) along with cognitive measures at visit 1 and 2, the random forest model predicted GEC at visit 3 with high accuracy (*r* = 0.61, *p* = 1.6e-09 between the true and predicted scores). Out-of-Bag (OOB) variable importance analysis revealed the top predictors of the model in clusters localized in the prefrontal and parietal cortices. Note, GEC = Global executive composite, OOB = Out-of-Bag.

**Figure 7.**
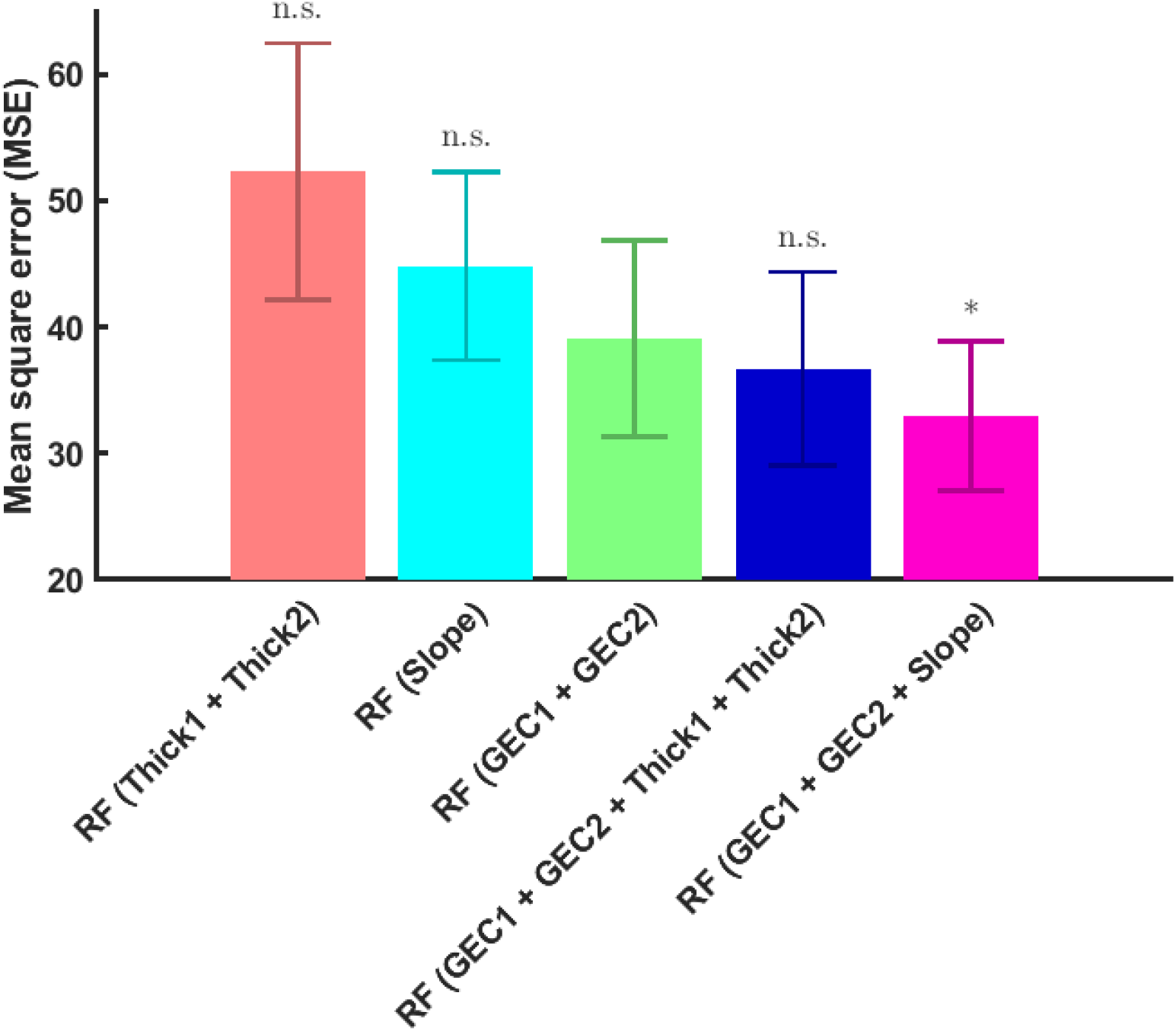
Comparison of prediction models of cognition at visit 3 using static and dynamic brain measures. Hypothesis testing was done with respect to the predictions of the random forest (RF) model using GEC at visit 1 and 2 (the baseline model). The prediction model performed poorly when demographic data (age and sex) or only brain measures were used as input data. Using static brain measures improved the prediction but was not significantly different from the baseline (*p* = 0.193), while using dynamic brain measures significantly improved the prediction (*p* = 0.028). Note, MSE = mean square error, * denotes *p* < 0.05

**Table 1.** Mean square errors (MSEs) of Random Forest (RF) models for predicting GEC_t3_ based on static and dynamic brain measures. Hypothesis testing compared MSEs of other models to RF model (GEC_t1_ + GEC_t2_), which we considered to be our baseline model. Note, RF = Random Forest, GEC = Global Executive Composite, MSE = mean square error, MAE = mean absolute error.

| **Model (Input data)** | **MSE** | **MAE** | ***p*** |
| --- | --- | --- | --- |
| RF model (Age + Sex) | 52.81 | 5.67 | 0.980 |
| RF model (Thick<sub>t1</sub> + Thick<sub>t2</sub>) | 52.27 | 5.70 | 0.981 |
| RF model (Slope) | 44.82 | 5.38 | 0.859 |
| RF model (GEC<sub>t1</sub> + GEC<sub>t2</sub>) | 39.07 | 4.72 | - |
| RF model (GEC<sub>t1</sub> + GEC<sub>t2</sub> + Thick<sub>t1</sub>) | 36.71 | 4.61 | 0.189 |
| RF model (GEC<sub>t1</sub> + GEC<sub>t2</sub> + Thick<sub>t2</sub>) | 37.29 | 4.67 | 0.244 |
| RF model (GEC<sub>t1</sub> + GEC<sub>t2</sub> + Thick<sub>t1</sub> + Thick<sub>t2</sub>) | 36.69 | 4.59 | 0.193 |
| RF model (GEC<sub>t1</sub> + GEC<sub>t2</sub> + Slope) | 32.97 | 4.51 | **0.028** |

## Discussion

In this paper, using longitudinal MRI data from 82 participants (scanned 3 times at ~2 years apart with concurrent measures of cognition), we built prediction models of future cognition (i.e. executive function at visit 3) using baseline developmental changes in cortical anatomy (measured as slope of the change in cortical thickness from visit 1 to 2). Our random forest model predicted future cognition (Global executive composite, an indicator of executive function at visit 3) with high accuracy (*r* = 0.61, *p* = 1.6e-09 between the true and predicted scores). The best predictors of the model were localized in clusters localized in the prefrontal and parietal cortices, regions that have been consistently shown associated with measures of executive function Comparative analysis revealed that the model using dynamic brain measures (but not static brain measures) significantly improved the prediction compared to the model which used only baseline cognitive measures (at visit 1 and 2).

The quest for understanding the neurobiology of cognitive development has relied on univariate methods that have revealed relationships between brain and cognitive measures (38–42). In line with previous studies, our univariate analysis also showed association of brain measures and future cognition (**Figure 2, 3**). Interestingly, a significant association of future cognition was observed with dynamic brain measures (**Figure 3**) but not with the static brain measures (**Figure 2**), indicating the relative importance of dynamic over static brain measures in the understanding of cognitive development. Although the univariate method elucidated the relationship between dynamic brain measures and future cognition, the full scope of dynamic brain measures in understanding cognitive development lies in predictive modeling of long-term cognitive outcomes based on baseline dynamic brain measures (26, 27).

With this motivation, we built prediction models (random forest models) of future cognition (Global executive composite, GEC at visit 3) using baseline brain and cognitive measures (at visit 1 and 2) as input data. Our findings revealed a highly accurate prediction model (*r* = 0.61, *p* = 1.6e-09 between true and predicted scores) using baseline dynamic brain measures (slope of change of cortical thickness from visit 1 to 2) and baseline cognitive measures (GEC at visit 1 and 2) (**Figure 6**). Not surprisingly, baseline cognitive measures (GEC at visit 1 and 2) in addition to distinct brain regions were observed as the top predictors of the model (**Figure 6B**). This finding was expected as the cognitive measure at visit 3 significantly correlated with that of visit 1 (*r* = 0.48, *p* < 0.001) as well as that of visit 2 (*r* = 0.56, *p* < 0.001). Findings of our comparative analyses further showed the importance of baseline cognitive measures in the prediction of future cognition. Prediction models performed worse if only brain measures (without the baseline cognitive measures) were used as input (**Table 1**, **Figure 7**). The models improved in performance when brain measures along with the baseline cognitive measures were used as input. However, it was inclusion of the dynamic (and not the static) brain measures that significantly improved model performance (**Table 1**, **Figure 7**), suggesting the relative importance of dynamic over static brain measures in predicting future cognition. Our findings can be interpreted in the light of previous trajectory-based studies (28, 43–48). Brain development comprises of sequence of events that continuously shape the structure and function of the brain (49–56). More importantly, trajectories of brain development have been shown to be critical for cognitive maturation (28, 43, 57), with deviations in normative trajectories linked with neurodevelopmental disorders (45, 58, 59). In light of this, it is possible that prediction models of cognition which incorporate developmental changes in brain measures would perform better than those with static brain measures.

In terms of brain regions, top predictors of the model were observed in clusters localized in the prefrontal and parietal cortices (**Figure 6B**), regions that have been consistently shown associated with measures of executive function (42, 60, 61). A systematic review of different clinical populations observed the frontal, parietal and cerebellar lobes as consistent neuroimaging correlates of executive function (61). Brain regions identified in our study are also part of the frontoparietal network (FPN), a flexible system that supports cognitive control (62–65). Interestingly, one meta-analysis of neuroimaging studies has shown that the frontoparietal network supporting cognitive control also supports a broad range of executive functions (29). Taken together, in consistent with previous studies, our findings highlight the critical role of a frontoparietal network in the maturation of executive function.

Some limitations of our study need to be mentioned. First, our study sample comprised of longitudinal neuroimaging data of 82 individuals (with a total of 246 scans) which is a relatively small sample size for brain-cognition association studies. Small-sample sized studies may lead to increased statistical errors such as false negatives, inflation undermining reproducibility (66). However, in spite of the relatively small sample size, our study utilized machine-learning based predictive modeling which has been shown to improve reproducibility of brain-cognition associations (66). Future studies will use large-scale, longitudinal data such as the Adolescent Brain Cognitive Development, ABCD database (67) to further validate and improve our methodology. Second, using principal component analysis, we reduced the high-dimensional neuroimaging data (81,924 vertices) to 14 principal components for model input. This raised the question of the optimal number of principal components. However, our supplementary analysis revealed that the performance of the prediction model stabilized after the inclusion of principal component 5 (**Supplementary Figure S1**) and remained stable thereafter, indicating the stability of our model. Lastly, we showed the scope of our prediction model using global executive composite, GEC (an indicator of executive function). Future studies will investigate other cognitive measures to determine the extent to which the findings of the present study can be generalized to other measures of cognitive function.

In conclusion, findings of our study revealed that dynamic brain measures (and not static brain measures) are important for predicting future cognition. Our findings also highlighted the scope of machine-learning based prediction models (as compared to traditional univariate methods) in understanding the neurobiology of cognitive development.

## Acknowledgements

This work was supported by The Azrieli Neurodevelopmental Research Program in partnership with Brain Canada Multi-Investigator Research Initiative (MIRI) grant to ACE. Linda Booij was supported by a chercheur boursier senior award from the Fonds de Recherche du Québec Santé. JT acknowledges support by the Academy of Finland (grant no 316258 to JT).

---

**Supplementary Figure 1:**
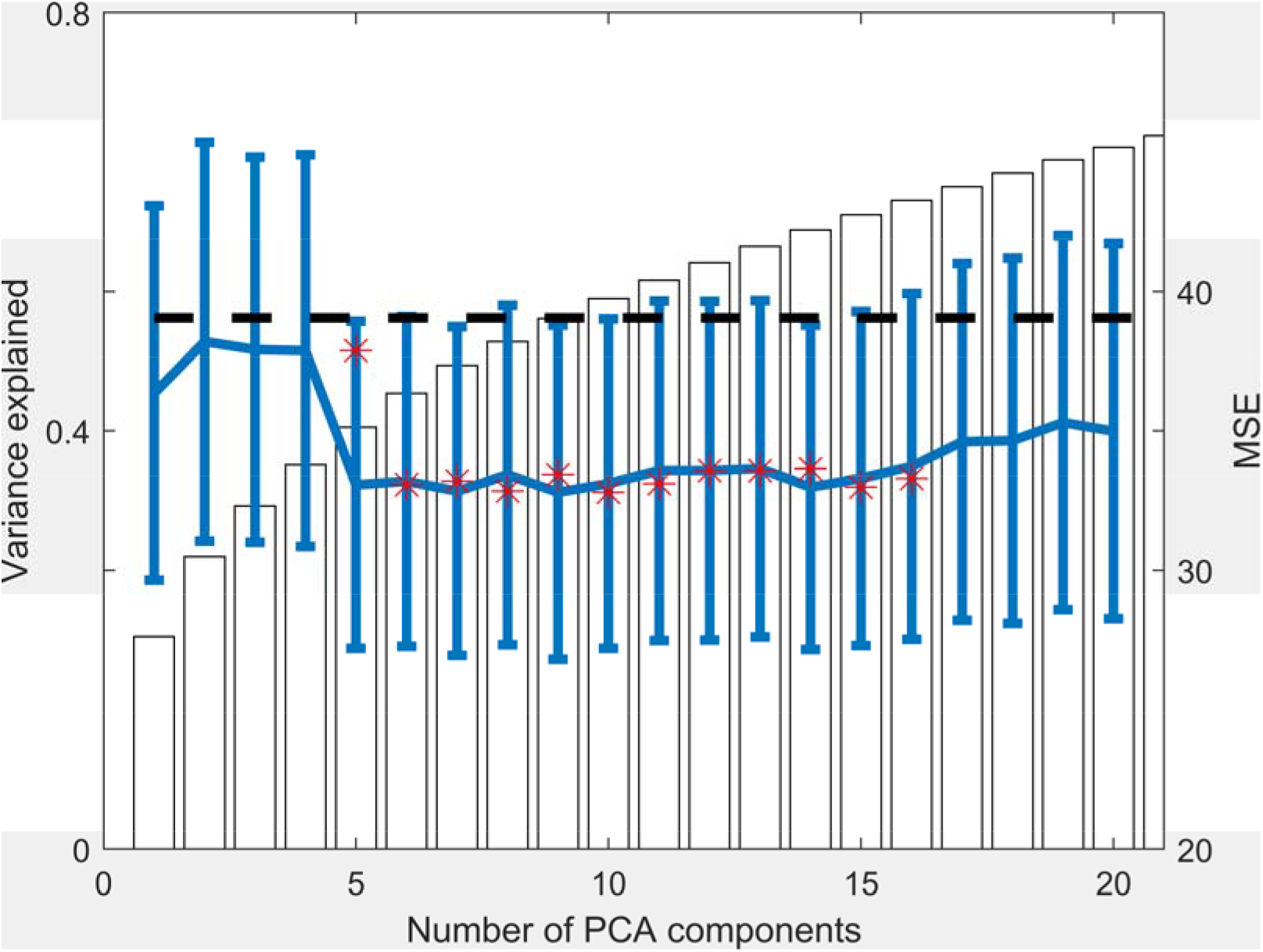
Model performance vs number of PCA components. The performance of the prediction model (as indexed by MSE) stabilized after inclusion of principal component 5 and remained stable thereafter, indicating the stability of our model. Note, PCA = principal component analysis, MSE = mean square error.

